# Single-Cell Mapping of Malignant Signaling Networks Guides Drug Combinations

**DOI:** 10.64898/2026.08.12.744506

**Authors:** Bengi Ruken Yavuz, Hyunbum Jang, Ruth Nussinov

## Abstract

Single-cell breast cancer atlases reveal malignant, immune, and stromal diversity; however, how the recurrent signaling pathways drive untreated malignant-cell states and could inform combination therapy remains unclear. Here, we analyzed 15,753 malignant cells from untreated primary breast tumors using a cell-resolved network framework. Individual transcriptomes were projected onto a protein–protein interaction network, partitioned into Leiden communities, and annotated by pathway enrichment. Pathway recurrence was evaluated against matched null models preserving community size, protein-network degree, and gene detection rate. Before null correction, recurrent pathways included PI3K/AKT, MAPK, JAK/STAT, and HIF-1 (hypoxia-inducible factor 1) signaling. After correction, HIF-1 emerged as the dominant recurrent signal across patients, indicating convergence of diverse upstream pathways on a shared hypoxia- and stress-adaptive malignant-cell program. The recurrent JAK/STAT, cAMP, glucagon, oxytocin, and thyroid hormone signaling suggest inflammatory, metabolic, and endocrine crosstalk. These findings support rational drug combinations targeting HIF-1 together with upstream PI3K/AKT/mTOR, MAPK, or JAK/STAT signaling.

## Introduction

Here, we identify recurrent intracellular signaling pathways with translational relevance, aiming to determine which signaling programs recur across malignant cells more often than expected. Knowledge of such recurrent pathways would enable clinicians to tailor optimal drug combinations which we suggest should be preferentially based on recurrent protein targets. Inhibiting these parallel or complementary pathways is expected to reduce the oncogenic signaling strength. Within this framework, we identify HIF-1 signaling as a convergent malignant-cell program, leading us to propose targeting HIF-1 pathway in combination with upstream growth-factor or cytokine-associated pathways, including PI3K/AKT/mTOR, Ras/MAPK, and JAK/STAT3.

Recent single-cell studies have mapped the cellular architecture of breast tumors across patients and clinical subtypes^1–3^. These analyses have linked malignant cell programs to their immune and stromal microenvironments and increasingly to spatial organization in tissues^1,4,5^. However, the recurrent intracellular signaling pathways that drive malignant cell states remain poorly characterized^6,7^. Single-cell transcriptomic data are sparse, noisy, and high dimensional. Network-based methods can identify coordinated signaling modules and prioritize candidate drivers of malignant progression and therapeutic resistance^6,8^.

Atlas-scale and cross-cohort analyses have linked malignant epithelial heterogeneity and tumor-immune system interactions to tumor subtype and therapy response^1,9^. Studies of primary and metastatic ER-positive breast cancer have resolved malignant and microenvironmental states at single-cell resolution^3^. Matched transcriptomic and epigenomic profiling have also begun to reveal their regulatory mechanisms^6,7^. More focused studies have shown that malignant cell states vary with estrogen response, metastatic dissemination, platinum resistance, circulating epithelial populations, and metastatic niche remodeling^10–14^.

This gap is important because malignant phenotypes rarely arise from isolated proteins alone^15–19^. Instead, they emerge from coordinated signaling networks that integrate growth-factor signaling, stress adaptation, metabolism, inflammation, survival, plasticity, and therapeutic or immune resistance^20–22^. Canonical oncogenic pathways such as PI3K/AKT/mTOR, Ras/MAPK, and JAK/STAT3 are central to breast cancer biology^23,24^. However, these pathways are highly connected and shared across cellular processes. Their frequent enrichment in single-cell pathway analyses can be difficult to distinguish from background expectations without an appropriate null model.

Network-based analysis provides a natural framework for addressing this problem. Rather than considering proteins in isolation, it places each cell’s transcriptome within a known molecular interaction network. This approach can identify connected protein groups that form recurrent pathway modules^25,26^. It is particularly useful for single-cell RNA-seq data where data are sparse but coordinated signals across modules can be informative^6,8,25^. Prior studies have uncovered immune-infiltration-associated hub genes, myeloid or macrophage-tumor signaling, CAF-mediated immune suppression, *SPP1*-*CD44* interactions, and therapy-induced remodeling of the triple-negative breast cancer (TNBC) immune microenvironment^27–30^. Although these studies have provided important insights into tumor–microenvironment interactions, they have not systematically identified recurrent intracellular signaling modules that drive malignant-cell states across patients.

Parallel computational efforts have increasingly translated single-cell transcriptomic data into therapeutic hypotheses by ranking drugs and drug combinations from patient-specific cellular states. Methods such as comboSC, RSDP, scTherapy, sc2MeNetDrug, and *retriever* have used single-cell transcriptomic data to identify personalized drug combinations, predict resistance-reversing therapies, infer cell–cell signaling perturbations, and integrate drug-response profiles with tumor atlases^31–35^. However, these methods generally prioritize therapeutic interventions from observed transcriptional states. Consequently, they overlook the intracellular signaling modules that are shared across malignant cells and patients after accounting for network and statistical background^36–38^. Consistent with this target-centered perspective, our earlier network-informed study showed that rational combination therapy should begin by identifying biologically meaningful co-targets before narrowing the selection to specific drug pairs^39,40^.

To address this gap, we applied a cell-resolved network framework to malignant cells from untreated primary breast tumors. We asked which intracellular signaling pathways recur across these cells more often than expected. Rather than relying on gene expression or pathway frequency alone, we mapped each malignant cell’s transcriptome onto a protein-protein interaction network and identified its highest-scoring network community. We then annotated the pathways represented in these cell-specific communities and compared their recurrence with matched null models that accounted for community size, protein-network connectivity, and gene detection rate. This approach distinguished broadly enriched canonical pathways from signals that recurred beyond expected network and detection biases. From a translational perspective, canonical upstream pathways such as PI3K/AKT/mTOR and Ras/MAPK remain biologically important. Our framework evaluates whether recurrent signaling through these pathways converges on more specific malignant-cell programs that can guide follow-up testing. In our analysis, HIF-1 emerged as such a convergent pathway. This calls for testing targeting HIF-1 directed pathway in combination with upstream growth-factor or cytokine-associated pathways, including PI3K/AKT/mTOR, Ras/MAPK, and JAK/STAT3.

## Results

### Curating an untreated malignant-cell cohort for network-guided pathway prioritization

We screened the Curated Cancer Cell Atlas (3CA)^41^ for breast cancer single-cell RNA-seq cohorts suitable for malignant cell network analysis. To minimize therapy-related transcriptional effects and cross-study technical batch effects, we selected a single primary breast cancer cohort with malignant-cell annotations, treatment-status data, and representation of the major clinical subtypes. On this basis, our downstream analysis focused on the Wu et al. 2021 cohort^1^ was selected for downstream analysis.

The selected cohort included 26 primary breast tumors: 11 estrogen receptor-positive (ER-positive), 5 HER2-positive, and 10 TNBCs. Across the full dataset, 100,064 cells were annotated across 13 major cell types, including malignant, immune, stromal, endothelial, and other epithelial populations (Figure S1a). T cells were the most abundant type, with 33,368 cells, followed by malignant cells, with 24,489 cells. Additional cell populations included endothelial cells, fibroblasts, macrophages, pericytes, plasmablasts, B cells, monocytes, NK cells, dendritic cells, and myeloid cells. Their counts ranged from 7,605 endothelial cells to 463 myeloid cells. Per-sample analysis further showed variation in both total cell number and cellular composition across tumors (Figure S1b).

We then restricted the analysis to malignant cells from untreated primary tumors to create the discovery dataset for intracellular network analysis (Figure 1). After preprocessing, quality filtering, and treatment-status restriction, the final dataset contained 15,753 malignant cells from 14 untreated primary breast tumors. All samples were invasive carcinomas from female patients, with a median age of 56.5 years. The final cohort included all three major clinical subtypes: seven ER-positive, five TNBC, and two HER2-positive tumors. Most tumors were invasive ductal carcinomas, with two invasive lobular carcinomas, and all cases were intermediate or high grade. Most samples with recorded nodal status were node-positive, and notable cases included one BRCA2-mutant TNBC and two basal-phenotype TNBCs. Thus, the curated dataset captured untreated malignant cells across clinically relevant breast cancer subtypes and higher-risk disease features. This cohort was then used for cell-specific network reconstruction and pathway-level analysis. As depicted in Figure 1, malignant-cell transcriptomes were mapped onto a reference protein-protein interaction network, cell-specific subnetworks were reconstructed, network communities were identified and enriched signaling pathways were annotated for downstream prioritization. This workflow provided a bridge from single-cell malignant-cell states to network-guided pathway hypotheses rather than relying on cell-type composition or differential expression alone.

**Figure 1.**
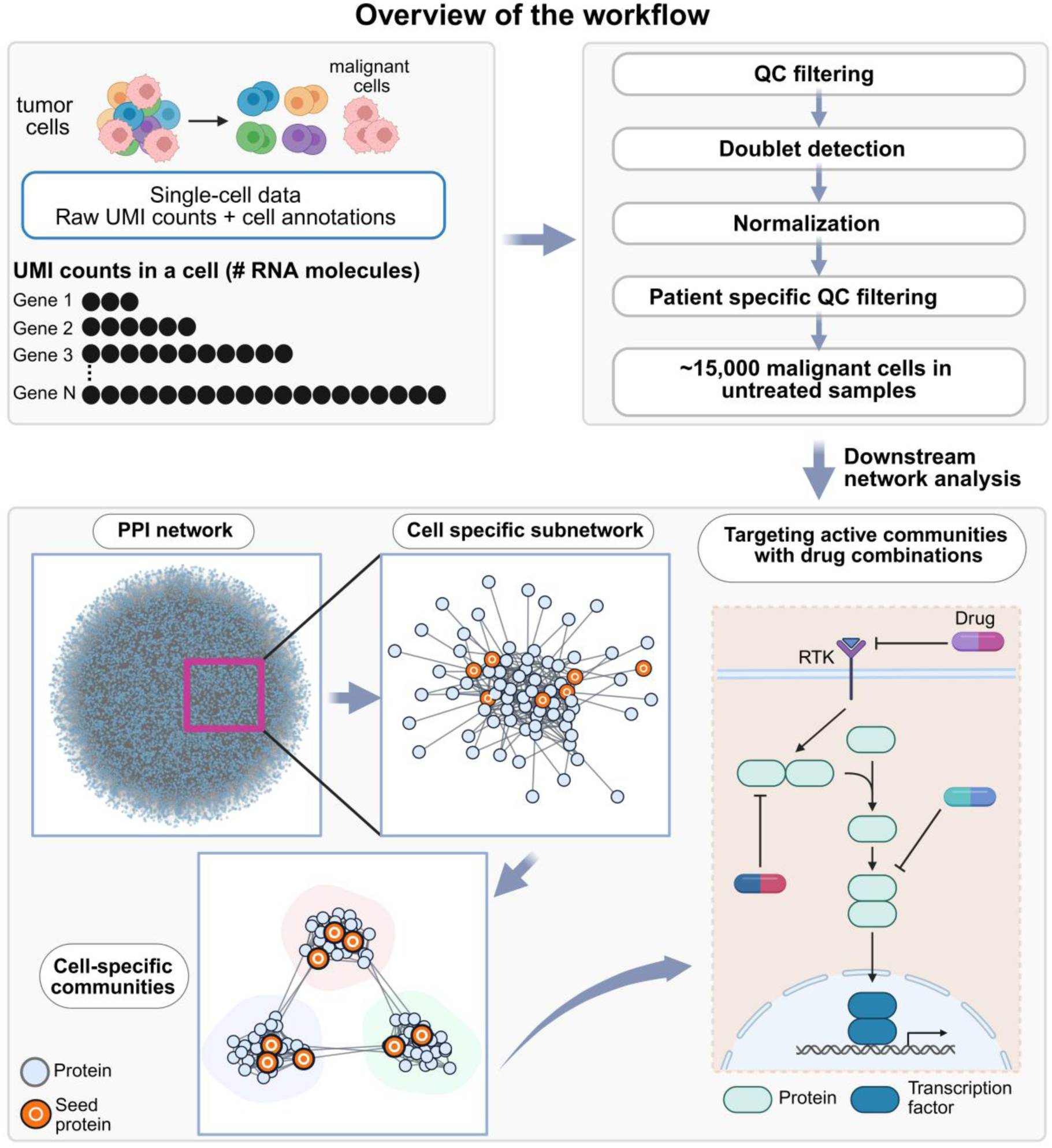
Single-cell preprocessing and cell-specific network-analysis workflow. The workflow proceeds from raw single-cell RNAseq data to filtered malignant-cell profiles and then to cell-resolved network reconstruction. In the upper panels, raw single-cell UMI (unique molecular identifier) count data and cell annotations from breast tumor samples are processed through sequential quality-control steps. These include removal of low-quality cells and rare genes, detection of doublets and retention of predicted singlets, normalization, control of sample-specific quality, and selection of malignant cells from untreated samples. This preprocessing for downstream analysis yielded 15,753 malignant cells across 14 untreated samples. Log-normalized expression values derived from the filtered UMI count matrix were then used for pathway scoring and network/module analysis. In the lower panels, each malignant-cell transcriptome is mapped onto the HIPPIE protein–protein interaction network to reconstruct a cell-specific subnetwork using PageRank, followed by Leiden community detection and signaling-pathway enrichment. The resulting cell-specific highest scoring Leiden communities and enriched pathways are used to prioritize highest-scoring signaling pathways and nominate drug-combination strategies after matched null correction.

After defining the untreated malignant-cell dataset, we examined its patient-level composition and the distribution of network-derived co-targeting candidates (Figure 2). These candidates were selected using our previously described framework, which prioritizes proteins with high betweenness centrality as connector nodes in reconstructed protein–protein interaction subnetworks^39^. The framework proposed co-targeting connector proteins that link alternative, redundant, parallel, or compensatory signaling routes. This strategy aims to disrupt oncogenic signaling and counter therapeutic resistance. Here, candidate proteins are labeled by their corresponding gene symbols.

**Figure 2.**
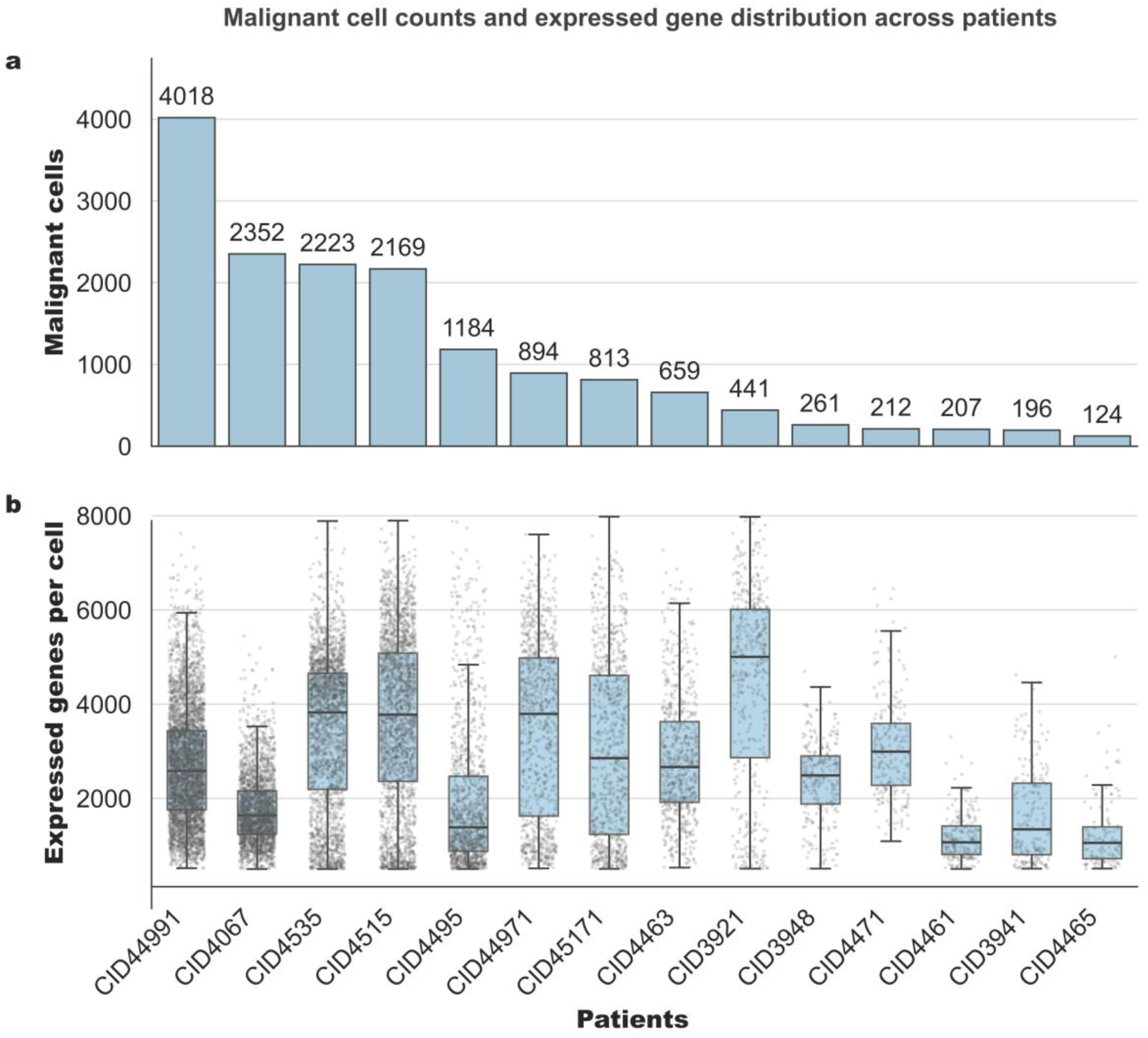
Patient-level malignant-cell counts and gene-detection distributions in untreated samples. **(a)** Number of malignant untreated cells retained for each patient after preprocessing. Each bar represents one patient with the malignant-cell count shown above the bar. Patients are ordered from highest to lowest malignant-cell count, from CID44991, which contributed the most cells, to CID4465, which contributed the fewest. Together, the bars represent the 15,753 malignant cells in the final dataset. **(b)** Distribution of the number of genes detected per malignant cell for the same patients, shown in the same order as in panel a. Each point represents one cell, and the boxplots summarize the within-patient distributions. Together, the panels show differences in patient contributions to the final dataset and variation in per-cell gene detection across patients.

Cell representation varied substantially across patients, ranging from 4,018 malignant cells in CID44991 to 124 malignant cells in CID4465 in the curated dataset (Figure 2a). The number of detected genes per cell varied across patients in both median and distributional spread, indicating differences in gene detection rates (Figure 2b). We then summarized, for each patient, the number of malignant cells associated with each co-targeting candidate retained in the cell-specific network analysis. The heatmap revealed patient-level variation in co-targeting candidate counts, with higher absolute counts generally observed in samples containing more malignant cells (Figure S2). The retained candidates include *EGFR*, *ERBB2*, *ERBB3*, *FGFR2*, *FGFR3*, *IGF1R*, and *GRB2*, Ras/MAPK pathway members including *ARAF*, *BRAF*, and *RAF1*, PI3K/AKT pathway members such as *AKT1* and *GSK3B*, and transcription factors or regulatory genes including *ESR1*, *MYC*, *MDM2*, and *STAT3*.

Together, these analyses revealed patient-level heterogeneity in malignant-cell representation, per-cell gene detection, and the distribution of network-derived co-targeting candidates. They provided the basis for reconstructing malignant-cell-specific subnetworks, identifying the highest-scoring Leiden community in each cell, and annotating its signaling pathways for recurrence analysis.

### Cell-resolved network propagation prioritizes malignant signaling-associated intracellular modules

To translate individual-cell expression profiles into interpretable signaling pathways, we reconstructed a protein–protein interaction subnetwork for each untreated malignant cell. Each transcriptome was mapped onto the HIPPIE^42^ protein-protein interaction (PPI) network using personalized PageRank. This algorithm extends standard PageRank by biasing the random walk toward predefined seed nodes. For each cell, the top 20% of detected genes ranked by raw UMI count served as cell-specific seeds. We also included 53 curated co-targeting candidate genes from our previous study study^39,43^ and assigned them fivefold greater weight. This weighting directed network propagation toward signaling-relevant regions of each cell-specific network.

Leiden community detection partitioned each cell-specific subnetwork into topology-defined intracellular modules. Each module contained genes detected in that cell whose encoded proteins formed a relatively connected neighborhood in the HIPPIE PPI network. We used these communities as candidate signaling modules for downstream enrichment analysis. Across 15,753 untreated malignant cells, this process yielded 104,046 cell–community entries (Supplementary Data 1). Within each cell, communities were ranked according to an expression-based community activity score, and the highest-scoring community selected for downstream analysis. We refer to this community as the dominant transcriptionally represented module. Here, “dominant” indicates the greatest expression-weighted representation in the reconstructed network and does not imply functional pathway activation.

The number of Leiden communities detected per cell was narrowly distributed across the untreated malignant-cell dataset (Figure S3a). Most cells contained seven communities (63.4%, n = 9,985), followed by six communities (33.5%, n = 5,275). Smaller fractions of cells contained five communities (3.1%, n = 481) or eight communities (0.1%, n = 12). Together, these groups accounted for all 15,753 untreated malignant cells. This consistency provided a common basis for selecting each cell’s highest-scoring module and conducting downstream pathway-enrichment analysis.

In contrast to the narrow range of Leiden community counts per cell, co-targeting candidates were sparsely and unevenly distributed across the resulting communities (Figure S3b). Most communities contained no co-targeting candidate (69.1%), whereas 13.6% contained one. Communities containing two, three, or four candidates accounted for 5.8%, 2.7%, and 1.9% of communities, respectively, whereas a smaller subset contained five or more candidates (7%). Thus, co-targeting candidates were concentrated in a minority of topology-defined modules rather than being uniformly distributed across the reconstructed subnetworks.

Communities without predefined co-targeting candidates likely represent expressed network modules outside the candidate set. Candidate-containing communities marked topology-defined neighborhoods containing one or more connector proteins. Communities with multiple candidates may indicate convergence of potentially targetable connectors within the same module. This distribution provides a practical basis for generating combination hypotheses. Candidates within the same community may support same-module or same-pathway co-targeting, whereas candidates in distinct but connected communities may indicate parallel or compensatory signaling pathways. Thus, candidate distribution links cell-specific network structure to experimentally testable combination strategies.

Community activity scores varied across cell-specific Leiden communities, reflecting variation in the aggregate transcriptional representation of topology-defined modules (Figure S4). Within each malignant cell communities were ranked by activity score. The highest-scoring community was retained as the dominant transcriptionally represented module. The activity scores of these selected communities differed across patients and varied among cells from the same patient (Figure S5). This indicates interpatient and intrapatient variation in the expression-weighted prominence of the selected network modules.

Co-targeting candidates were unevenly represented across Leiden communities and patients (Figure 3). In the all-community analysis, each candidate was counted in every Leiden community in which it occurred. The selected-community analysis considered only the highest-scoring community from each untreated malignant cell (Figure 3a). The bar lengths represent absolute community counts and provide a descriptive, rather than proportional, comparison of candidate occurrence.

**Figure 3.**
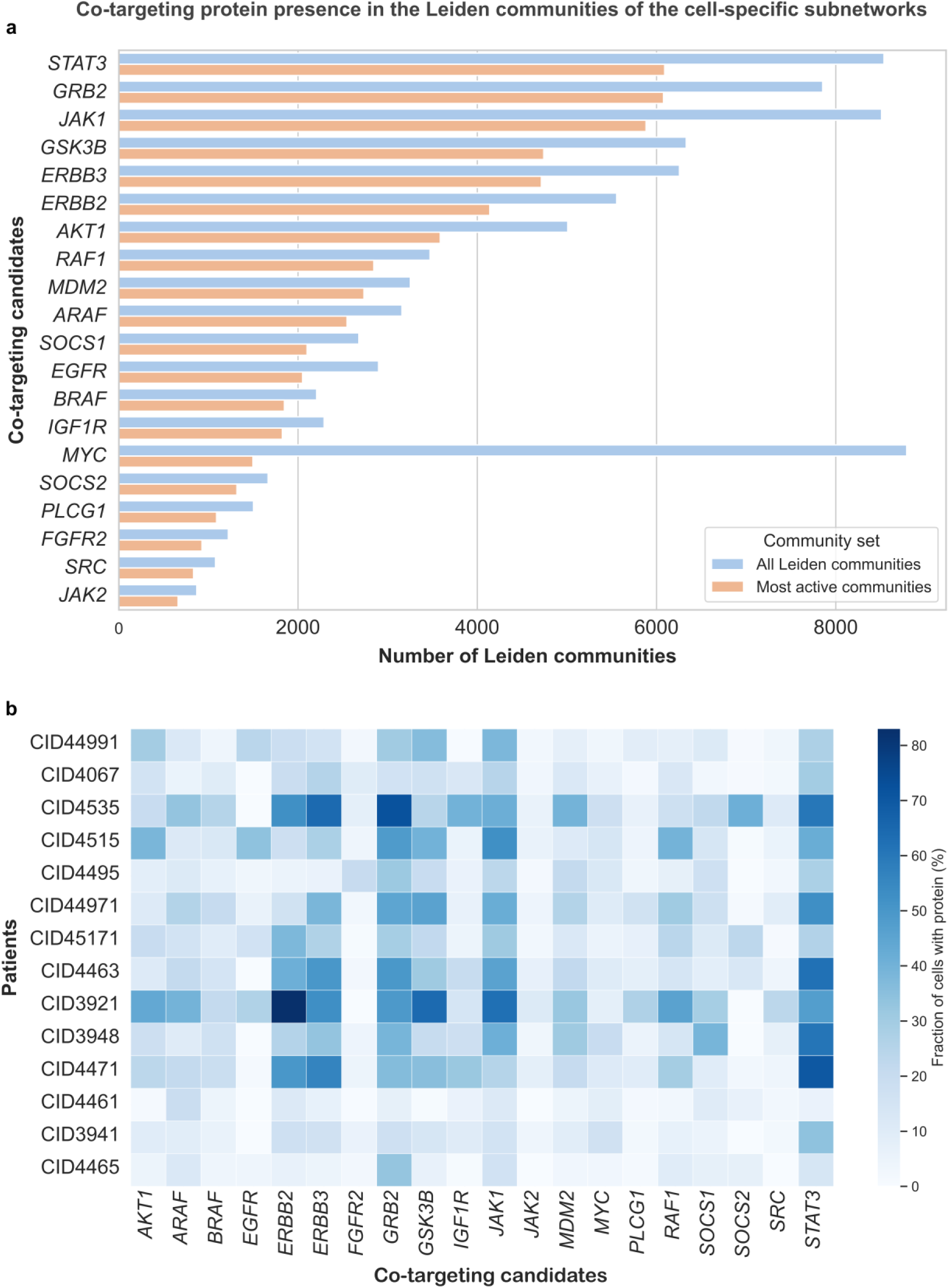
Representation of co-targeting candidates across Leiden communities and patients. Candidate proteins are labeled by their corresponding gene symbols. **(a)** Horizontal bar plot showing the number of Leiden communities containing each candidate. Blue bars include all communities detected across untreated malignant cells, whereas orange bars include only the highest-scoring community selected from each cell. Because one community was selected per cell, the orange bars also indicate the number of malignant cells whose highest-scoring module contained each candidate. Candidates represented in more selected communities may therefore be prioritized for follow-up co-targeting suggestions, particularly when complementary candidates occur within the same or connected modules. The bars show absolute counts and are intended for descriptive comparison. **(b)** Heatmap showing patient-level representation of co-targeting candidate proteins in the selected highest-scoring Leiden communities. Rows represent patients, columns represent co-targeting candidate proteins, and color intensity indicates the percentage of cells from each patient whose selected community contained the indicated candidate. The heatmap summarizes heterogeneity in candidate representation across patients and candidates.

*STAT3, GRB2, JAK1, GSK3B, ERBB3,* and *ERBB2* were represented in both community sets. *MYC* occurred frequently across all Leiden communities but was detected in fewer communities after the analysis was restricted to the highest-scoring community from each cell. At the patient level, candidate representation was measured as the percentage of cells whose selected community contained each candidate (Figure 3b). Together, these patterns provide a map of candidate-containing network modules for downstream pathway and co-targeting hypothesis generation.

### Recurrent HIF-1 enrichment links malignant-cell modules to hypoxia-adaptive and growth-factor signaling pathways

We first summarized KEGG^44^ signaling-pathway enrichment across the highest-scoring Leiden community in each untreated malignant cell before matched-null correction (Figure 4). This pre-null analysis recovered several expected breast cancer signaling pathways including HIF-1, and MAPK signaling, followed by prolactin, insulin, PI3K/AKT, sphingolipid, Rap1, thyroid hormone, and chemokine (Figure 4). The inset in Figure 4 further showed that most cells had a limited number of significant signaling pathways. The selected communities therefore showed focused pathway enrichment rather than broad, nonspecific enrichment across many signaling pathways.

**Figure 4.**
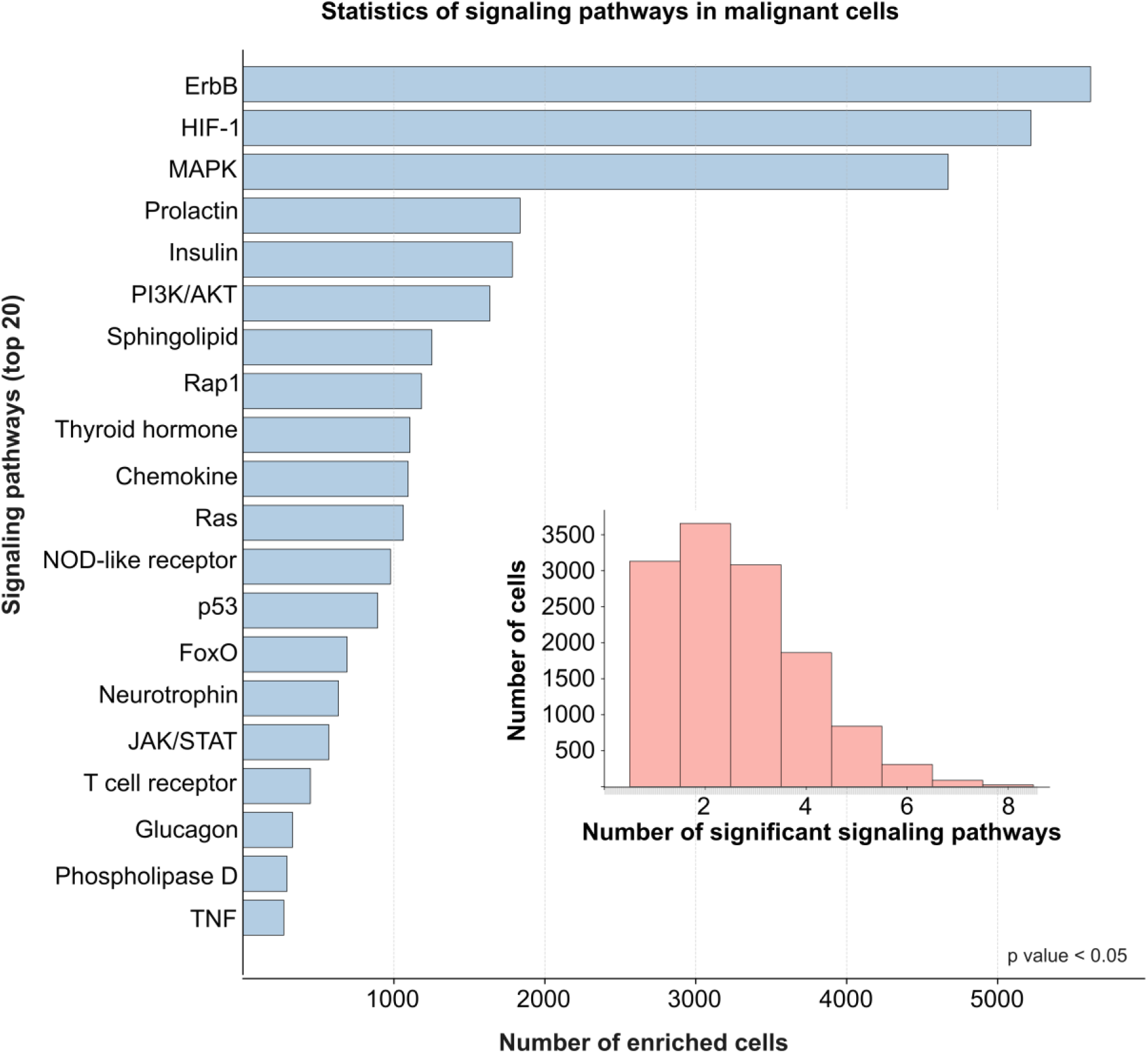
Recurrent signaling-related KEGG pathway enrichment before matched-null comparison. Horizontal bar plot showing the top recurrent signaling-related KEGG signaling pathways before matched-null correction. Bars indicate the number of malignant untreated cells whose highest-scoring Leiden community showed significant enrichment for proteins associated with each KEGG signaling pathway. Bar labels indicate the number of enriched cells for each pathway. The inset shows the distribution of the number of significant enriched signaling pathways per cell.

To identify pathways that recurred beyond background expectations, we compared the observed KEGG^44^ enrichment with matched-null distributions. For each patient–pathway pair, the observed enrichment fraction was the proportion of untreated malignant cells whose highest-scoring Leiden community was significantly enriched for that pathway by over-representation analysis. The null distribution was generated from 300 permutations. Each permutation preserved the cell-specific community gene-set size and matched genes by PPI degree and gene detection rate. A pathway was classified as recurrent after null adjustment only if it met the prespecified empirical q-value and Z-score thresholds and showed a positive observed-minus-null effect.

After matched-null correction, several KEGG signaling pathways remained enriched above background expectations in the highest-scoring Leiden communities (Figure 5). Among the displayed pathways, HIF-1 showed the largest median observed-minus-null difference, with a median Δ of 25.7% and a median standardized enrichment Z-score of 18.2. The matched-null baseline for HIF-1 signaling was also relatively high, indicating that HIF-1-associated proteins were frequently represented in the matched random sets. Nevertheless, observed HIF-1 enrichment remained substantially higher after accounting for community size, PPI degree, and gene detection rate. Rap1, JAK/STAT, sphingolipid, prolactin, Ras, and RIG-I-like receptor signaling also exceeded their matched-null medians. cAMP, thyroid hormone, oxytocin, phospholipase D, glucagon, and Wnt signaling showed smaller positive differences. Together, these findings identify a subset of KEGG signaling pathways that recurred in malignant-cell network modules more often than expected under the matched-null model.

**Figure 5.**
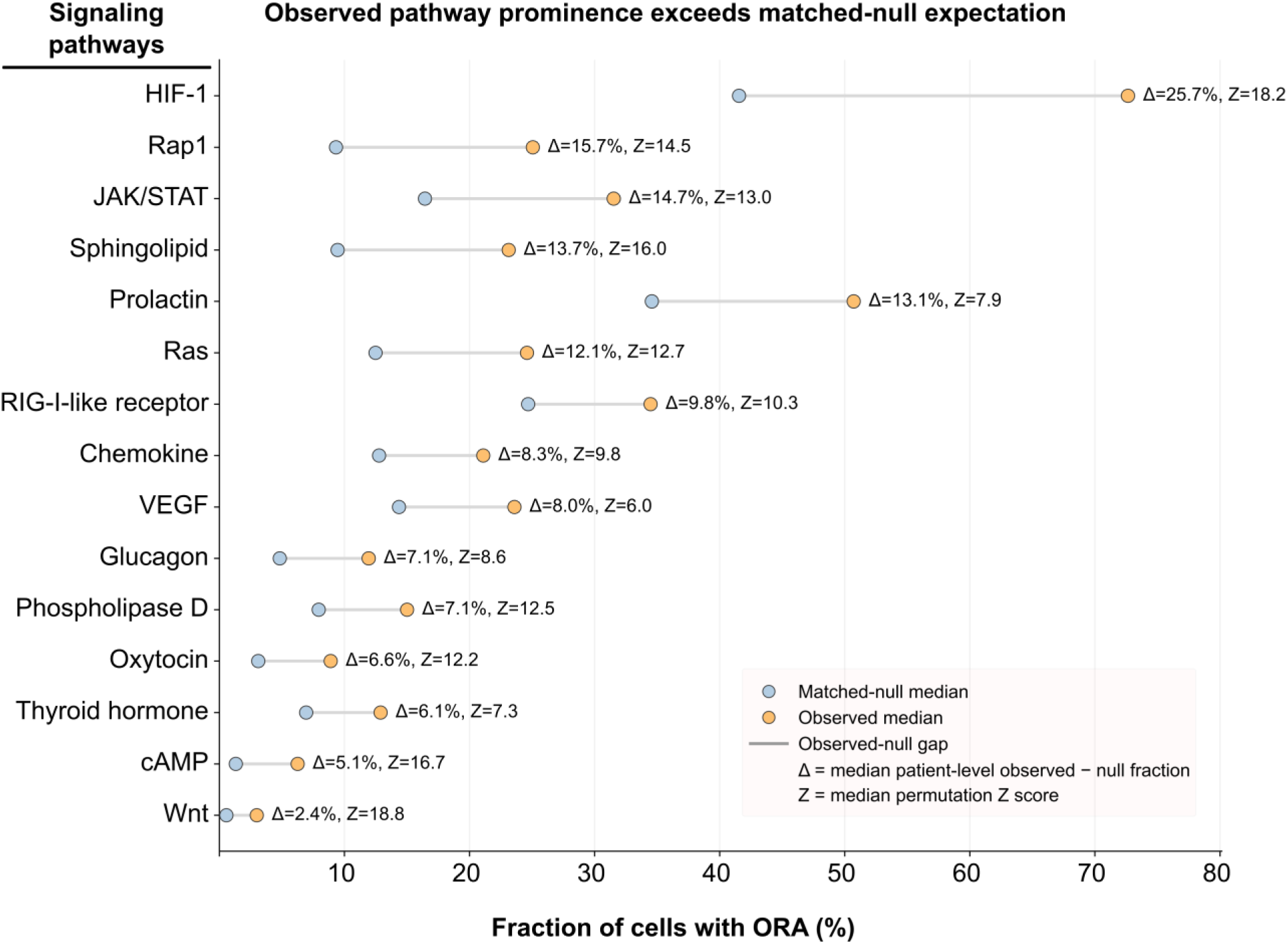
Observed frequencies of KEGG pathway enrichment compared with degree- and detectability-matched null expectations. Each row represents a KEGG signaling pathway identified as significantly more recurrent than expected under the matched null model. Orange and blue points indicate the median observed and matched-null fractions of cells with significant over-representation analysis (ORA) enrichment, respectively; gray segments show the observed-minus-null gap. Null permutations preserved community gene-set size and matched genes by PPI degree and gene detection rate. HIF-1 showed the largest excess over matched-null expectation (Δ = 25.7%, Z = 18.2), followed by Rap1, JAK/STAT, sphingolipid, prolactin, and Ras signaling. Additional immune, metabolic, endocrine, and stress-associated pathways, including RIG-I-like receptor, chemokine, VEGF, glucagon, oxytocin, thyroid hormone, cAMP, and Wnt signaling, also exceeded matched-null baselines.

At the patient level, HIF-1 signaling was the most recurrent null-adjusted KEGG pathway, with positive observed-minus-null effects, i.e., more of a specific biological activity than would be expected by random chance, detected in all 14 patients. Thyroid hormone and oxytocin signaling also showed broad positive effects, each detected in 13 patients, followed by glucagon, cAMP, and JAK/STAT signaling in 10, 9, and 8 patients, respectively. Representative HIF-1 comparisons showed observed enrichment fractions above matched-null expectations in CID3921 (0.816 vs. 0.589), CID4535 (0.801 vs. 0.544), CID4495 (0.791 vs. 0.343), and CID4515 (0.733 vs. 0.542). MAPK signaling was frequent before correction (Supplementary Data 2). Thus, matched-null testing distinguished pathways that were frequently enriched from those that recurred beyond background expectations. These findings identify HIF-1 as the most consistent null-adjusted pathway across patients, rather than a signal explained solely by its high pre-null enrichment frequency.

The recurrent enrichment of HIF-1 signaling suggests that the selected malignant-cell modules were associated with hypoxia- and stress-adaptive programs. This interpretation is consistent with the established role of HIF-1 in cellular oxygen sensing. Oxygen-dependent, VHL-mediated proteasomal degradation regulates HIF-α subunit stability^45,46^. HIF-1 is also implicated in cancer-associated transcriptional programs involving metabolic adaptation, angiogenesis, survival, invasion, apoptosis, and therapy resistance^47,48^. In breast cancer, HIF-related programs have been linked to metastatic progression^49^. A recent meta-analysis further associated high HIF-1α expression is with shorter overall and disease-free survival^50^.

To interpret the recurrent HIF-1 signal, we depicted a HIF-centered signaling schematic (Figure 6) that separates canonical oxygen-dependent regulation from cytokine- and growth-factor-mediated inputs. In the canonical oxygen-dependent pathway, newly synthesized HIF-1α is hydroxylated mainly by PHD enzymes, enabling VHL recognition and ubiquitin–proteasome degradation. In parallel, FIH limits HIF-1α transcriptional activity by reducing its ability to recruit p300/CBP. Therefore, under normoxia, HIF-1α protein remains unstable and HIF-dependent gene transcription is low^51–53^. Under hypoxia, reduced PHD and FIH activity stabilizes HIF-1α, allowing it to enter the nucleus, dimerize with HIF-1β/ARNT, recruit p300/CBP, bind hypoxia-response elements. This activates genes that promote tumor adaptation through angiogenesis, metabolic reprogramming, survival, invasion, immune escape, stemness, and drug resistance^51,54^.

**Figure 6.**
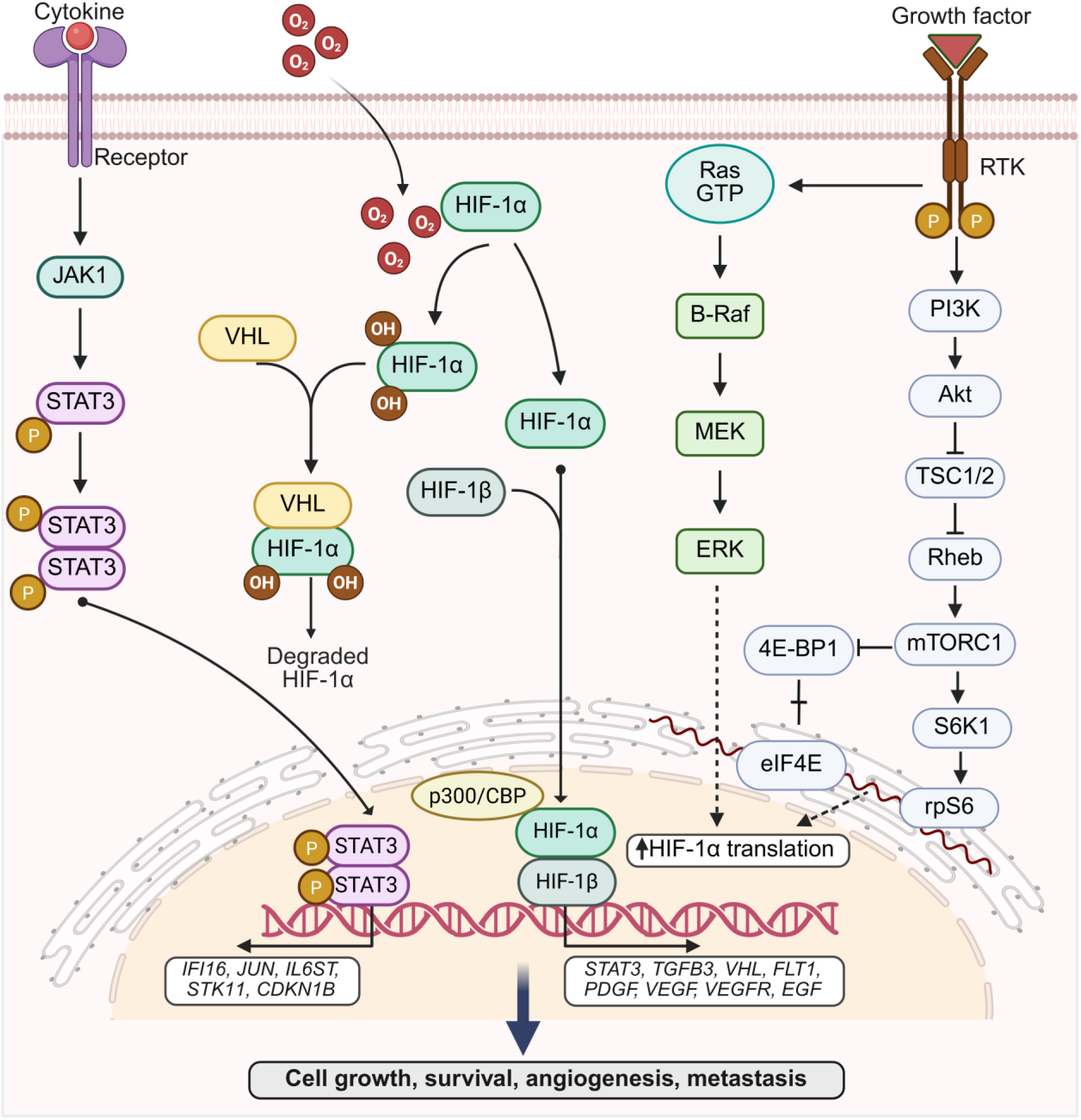
Schematic of upstream PI3K/AKT, Ras/MAPK, and JAK/STAT signaling converging on HIF-1α. Oxygen-dependent hydroxylation of HIF-1α, indicated by OH, promotes its association with VHL and subsequent degradation. Non-degraded HIF-1α translocates to the nucleus, dimerizes with HIF-1β, and acts with p300/CBP at DNA to regulate the target-genes. Growth factor binding activates RTK signaling and stimulates two downstream arms: in the PI3K arm, PI3K activates *AKT*; *AKT* inhibits *TSC1/2*; *TSC1/2* inhibition relieves repression of Rheb; and Rheb activates mTORC1. mTORC1 promotes the module shown by reducing 4E-BP1 mediated inhibition of eIF4E and activating the S6K1–rpS6 axis, thereby increasing HIF-1α translation. In the Ras/MAPK pathway, RTK signaling activates Ras-GTP, B-Raf, MEK, and ERK; ERK is shown as an indirect input that also promotes increased HIF-1α translation. In parallel, cytokine receptor signaling activates *JAK1* and phosphorylated *STAT3* dimers, which enter the nucleus and regulate the *STAT3* target genes. The *STAT3* and HIF-1 target genes include *IFI16, JUN, IL6ST, STK11, CDKN1B* (left rectangle) and *STAT3, TGFB3, VHL, FLT1, PDGF, VEGF, VEGFR, EGF* (right rectangle), respectively. Together, these signaling inputs support cell growth, survival, angiogenesis, and metastasis. Solid arrows indicate activation or directional signaling, solid arrows with a filled dot indicate translocation, solid arrows marked by a short transverse line indicate dissociation, blunt-ended lines indicate inhibition, dashed arrows indicate indirect regulation, P indicates phosphorylation, and OH indicates hydroxylation. Created with BioRender.com.

The schematic also shows how growth-factor receptor signaling converges on HIF-1α through PI3K/AKT/mTORC1 and Ras/MAPK pathways.

This regulation can occur independently of oxygen. Activated RTKs, including *EGFR* or *HER2*, stimulate PI3K/AKT/mTORC1 signaling. *AKT* inhibits *TSC1/2*, relieving repression of Rheb and activating mTORC1. mTORC1 then promotes HIF-1α synthesis through 4E-BP1/eIF4E and S6K1/rpS6 translational modules rather than by directly blocking VHL-mediated degradation^55,56^. In some contexts, PI3K/AKT increases HIF-1α translation under both normoxic and hypoxic conditions. This effect can be largely rapamycin-insensitive and mTOR-independent in standard serum culture conditions^57^. In invasive breast carcinoma, PI3K inhibition reduced hypoxia-induced HIF-1α levels, whereas active AKT1 restored them. AKT1 phosphorylation also correlated with HIF-1α expression in human breast tumors^58^.

The Ras/MAPK pathway provides a second growth-factor-associated input into HIF-1α regulation. RTK signaling activates Ras (by converting it to its GTP-bound state), B-Raf, MEK, and ERK^59^. ERK is represented in Figure 6 as an indirect input into HIF-1α translation and HIF-1 transcriptional activity. ERK/MAPK signaling can enhance HIF-1α function by increasing HIF-1α phosphorylation, nuclear retention, transcriptional activity, and cooperation with p300/CBP. ERK-dependent phosphorylation of HIF-1α at Ser641/Ser643 supports nuclear accumulation and transcriptional activity^60,61^. Oncogenic Ras can therefore activate HIF-1α output through both PI3K/AKT and Ras/MAPK pathways^52,62^.

The cytokine/JAK/STAT3 pathway provides another regulatory input into HIF-1 signaling. Cytokine receptors activate JAK1/JAK2, leading to *STAT3* phosphorylation, STAT3 dimerization, and nuclear translocation*. STAT3* can support the HIF-1 pathway in three ways: it can promote HIF1A transcription, stabilize HIF-1α in some cancer models by reducing VHL-mediated ubiquitination, and cooperate directly with HIF-1α at target-gene promoters^63–65^.

In MDA-MB-231 breast cancer and RCC4 renal carcinoma cells, *STAT3* binds HIF-1 target promoters, interacts with HIF-1α, and recruits p300/CBP and RNA polymerase II. These interactions promote hypoxia-dependent activation of HIF-1 target genes^66^. Dinarello et al. further showed that *STAT3* is required for full induction of a subset of hypoxia-responsive genes, including *VEGFA*, without affecting HIF-1α mRNA expression or HIF-1α protein stabilization. Instead, *STAT3* physically interacted with HIF-1α and was required for proper STAT3/HIF-1α-dependent transcriptional and physiological responses to hypoxia in cells and zebrafish^67^. EGF signaling can also connect *STAT3* to HIF-1α. In SW480 colorectal cancer cells, EGF induced *STAT3* phosphorylation and nuclear translocation, and phosphorylated *STAT3* was required for EGF-induced HIF-1α upregulation. This EGF–pSTAT3–HIF-1α axis promoted proliferation, migration-related behavior, and tumorigenesis^68^.

The HIF-centered schematic provides a biological framework for interpreting the null-adjusted enrichment results and translating recurrent pathway annotations into testable co-targeting hypotheses. It places HIF-1 at the intersection of oxygen-dependent VHL regulation, growth factor–driven PI3K/AKT/mTOR and Ras/MAPK signaling, and cytokine-associated JAK/STAT3 signaling. In breast cancer, HER2/PI3K/AKT/mTOR signaling can increase HIF-1α abundance and activity^55,58^. *STAT3* can cooperate with HIF-1α at target-gene promoters ^66^ and connect growth signaling to HIF-1/VEGF output^63^. Together, these mechanisms link the matched-null HIF-1 signal to hypoxia responses, oncogenic growth signaling, and *STAT3*-dependent transcription. This convergence provides a biological rationale for testing pathway-based combination strategies.

## Discussion

Our study extends single-cell breast cancer analysis beyond cell-state annotation by identifying recurrent intracellular signaling pathways in malignant cells. This framework reveals a key distinction between pathways that are frequently enriched and those that recur beyond background expectations. PI3K/AKT, HIF-1, MAPK, and JAK/STAT signaling were prominent before null correction. After accounting for community size, PPI degree, and gene detection rate, HIF-1 remained the most consistent recurrent pathway across patients. This persistence supports HIF-1 as a shared point of convergence in malignant cells from untreated breast tumors. Its network associations with PI3K/AKT, MAPK, and JAK/STAT further indicate that hypoxia-associated signaling may connect multiple oncogenic and stress-response pathways^24,69^.

HIF-1 links oxygen availability to transcriptional programs regulating glycolysis, angiogenesis, survival, invasion, stem-like phenotypes, immune modulation, and treatment resistance^47,51^. In breast cancer, high HIF-1α expression is also associated with shorter overall and disease-free survival^49,50^. HER2-driven signaling can increase HIF-1α protein abundance and *VEGF* expression through PI3K/AKT/mTOR pathway^55,58,70–72^. *STAT3* can also cooperate with HIF-1α at HIF-1 target gene promoters. In addition, IL-6/HIF-1 signaling has been linked to breast cancer stem cell-related phenotypes^66,67,73^.

PI3K/AKT/mTOR signaling regulates HIF-1α abundance and HIF-dependent transcription in breast cancer models. This relationship provides a biological rationale for co-targeting HIF-1 and PI3K/AKT/mTOR signaling^57,70^. *STAT3* can also cooperate with HIF-1α at hypoxia-responsive target genes. In triple-negative breast cancer, HIF-1α has been associated with phosphorylated *STAT3*^66,74^. When direct inhibition of the HIF axis is not feasible, dual perturbation of PI3K/AKT/mTOR and JAK/STAT3 signaling may offer a tractable strategy. The present study supports target-pathway prioritization over selection of specific drugs.

The recurrence of cAMP, glucagon, oxytocin, and thyroid hormone signaling highlights endocrine and metabolic regulation in malignant cell network states. Because these KEGG pathways share intracellular signaling components, their recurrence likely indicates convergence of endocrine-like, metabolic, growth-factor, and hypoxia-responsive signaling networks rather than direct hormonal stimulation alone^10,12^.

Recent therapeutic advances have expanded opportunities for genotype-guided targeting of PI3K and Ras/MAPK signaling^75–77^. In *PIK3CA*-mutant, HR-positive/HER2-negative advanced breast cancer, alpelisib established PI3Kα inhibition with endocrine therapy. Building on this, inavolisib improved outcomes when added to palbociclib and fulvestrant^78,79^. *KRAS* inhibition has also expanded from approved *KRAS*^G12C^ inhibitors to newer agents such as divarasib and olomorasib. Daraxonrasib (RMC-6236) more broadly targets GTP-bound mutant and wild-type RAS and has shown activity in RAS-mutant pancreatic cancer ^80–82^.

These advances provide combination of PI3Kα inhibition, preferably inavolisib or other clinically advanced PI3Kα inhibitors, with genotype-matched *KRAS*^G12C^ inhibitors for *KRAS*^G12C^-mutant tumors. In broader RAS-driven settings, daraxonrasib could serve as an alternative RAS-directed inhibitor. Because PI3K/AKT/mTOR and Ras/MAPK signaling can regulate HIF-1α abundance and hypoxia-responsive transcription, these combinations could test whether concurrent upstream inhibition reduces HIF-1-associated malignant-cell plasticity and stress adaptation.

Single-cell transcriptomics does not directly measure protein abundance, phosphorylation, or spatial organization. Nevertheless, our network-based analysis provides a framework for prioritizing recurrent signaling programs in malignant cells. Future perturbation studies should compare combined HIF-1 and PI3K/AKT/mTOR inhibition with combined HIF-1 and JAK/STAT3 inhibition, evaluating their respective effects on cell viability, apoptosis, glycolysis, angiogenesis, stem-like phenotypes, and therapy-resistant states.

Overall, our analysis identifies HIF-1 signaling as a recurrent and null-adjusted signaling pathway in untreated malignant breast cancer cells. This recurrence persists after matched-null correction for community size, PPI degree, and gene detection rate. This robustness argues against enrichment driven by pathway size or network topology and positions HIF-1 as a potential convergence point for PI3K/AKT/mTOR, RAS/MAPK, and JAK/STAT3 signaling. Together, these pathways may reinforce a shared hypoxic, metabolic, inflammatory, and stress-adaptive state across patients. Ultimately, this framework guides drug development by prioritizing pathways, helping evaluate how HIF-axis disruption interacts with targeted inhibitors—such as PI3K, RAS, or JAK—before selecting drug combinations.

## Methods

### Dataset selection and preprocessing

Breast cancer single-cell RNA-seq datasets in the Curated Cancer Cell Atlas were screened for human primary breast tumors with untreated samples and malignant-cell annotations. The Wu et al. 2021 cohort met these criteria and was selected for analysis^1^. The cohort comprised 26 breast tumor samples profiled using 10x Genomics platform. Treatment status was assigned from sample-level metadata. Nineteen samples were treatment-naïve (untreated), five were treated, and two lacked treatment-status information. Treated samples and those with missing treatment status were excluded. Downstream analyses were restricted to malignant cells from untreated samples.

Raw UMI counts were imported from a MatrixMarket file and transposed into AnnData objects, with cells as observations and genes as variables. Gene symbols were assigned from the accompanying annotation file, duplicate symbols were made unique, and cell-level metadata were added.

Quality-control metrics were calculated using Scanpy^83^, including the number of detected genes per cell, total UMI counts, and the percentage of counts assigned to mitochondrial genes. Mitochondrial genes were defined by MT-prefix. Cells with fewer than 200 detected genes or a mitochondrial-count fraction of at least 25% were excluded. Genes detected in fewer than five cells were also removed. These filters removed 3,049 genes but no cells, leaving 100,064 cells and 26,684 genes. Putative doublets were identified using Scrublet with an expected doublet rate of 0.06. Eight predicted doublets were removed, leaving 100,056 predicted singlet cells.

Expression matrices from predicted singlets were normalized to 10,000 counts per cell and log1p-transformed. We then applied an adaptive patient-level quality-control filter to account for differences in sequencing depth and gene-detection complexity. For each patient, the lower threshold for detected genes per cell was defined as the median minus 1.5 times the interquartile range, with a minimum threshold of one gene. Patients with fewer than 300 cells were excluded. The adaptive filter removed 76 cells, leaving 99,980 cells and 26,684 genes.

Malignant cells were selected using existing cell-type annotations. Cells were classified as malignant if the cell_type field contained “tumor” or “malignant.” This yielded 24,489 malignant cells. The final dataset used for network and pathway analyses contained 15,753 malignant untreated cells and 26,684 genes.

### Protein-protein interaction network construction and cell-specific network propagation

A reference PPI network was used from the HIPPIE interactome^42^. Each interaction between two gene symbols represented as an undirected edge with unit weight. The resulting graph served as a shared network backbone for all cells. Cell specificity was introduced through expression-derived personalization vectors used to compute personalized PageRank scores.

Each untreated malignant cell was mapped onto the reference PPI network using personalized PageRank^84,85^. For each cell, all network nodes were assigned a baseline personalization value of 1.0. Among the detected genes, the top 20% ranked by raw UMI count were selected. For selected genes present in the PPI network, baseline values were replaced with their cell-specific UMI counts.

A prespecified set of 53 co-targeting candidate proteins^39^ were then given fivefold greater wight. Candidate genes with UMI-derived values were multiplied by five, whereas the remaining candidate nodes were assigned a value of 5.0. Personalized PageRank was calculated with a damping factor of 0.85. This weighting prioritized the proteins encoded by these highly expressed genes in each cell while maintaining additional priority for co-targeting candidates.

### Leiden community detection and community activity scoring

Leiden community detection was performed independently on each cell-specific PPI subnetwork using leidenalg (https://github.com/vtraag/leidenalg) with a resolution parameter of 1.0. For subnetworks with at most one vertex or no edges, all available genes were assigned to community 0. Community activity score was calculated as the sum of normalized expression values for genes detected in the cell, defined by a raw UMI count greater than zero. Within each cell, communities were ranked by activity score, with ties resolved by the number of detected genes. The highest-scoring community was selected for pathway-enrichment analysis.

### Pathway enrichment analysis

Pathway enrichment analysis was performed on genes detected in the highest scoring Leiden community from each untreated malignant cell. Cells whose selected community contained fewer than 10 detected genes were excluded. Over-representation analysis was performed using Enrichr^86^ through gseapy.enrichr, with the KEGG 2021 human gene set library. The Enrichr cutoff was set to 1.0 to retain all enrichment results. The top 20 KEGG terms per cell were retained.

### Matched permutation testing of pathway prominence

Matched permutation testing was used to determine whether KEGG^44^ signaling pathways were enriched across malignant cells from each patient more often than expected under matched random gene sets.. For each patient, pathway prominence was defined as the number of cells in which a pathway achieved a nominal one-sided hypergeometric enrichment p-value of ≤0.05. Candidate pathways were ranked by their observed cell counts in the cell-level enrichment results. By default, analysis was restricted to signaling-related terms, and the top 25 pathways were retained.

Random gene sets were matched using two gene-level covariates: PPI degree and gene detection rate. PPI degree was calculated from the HIPPIE interactome. Gene detection rate was defined as the number of retained cell-specific gene sets containing each gene. Each covariate was divided into five quantile-based bins, producing 25 joint degree–detection bins. For each cell, a random gene set matching the size of the observed set was sampled to approximate its distribution across these joint bins.

For each patient–pathway pair, 300 matched permutations were performed. In each permutation, cell-specific random gene sets were generated, hypergeometric enrichment p-values were recalculated, and pathway prominence was recorded as the number of cells with P≤0.05^87,88^. The empirical one-sided p-value was calculated as

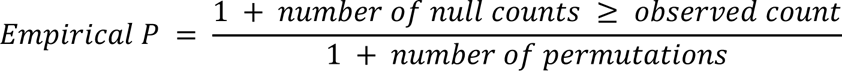

where the number of permutations is 300. Thus, the minimum attainable empirical p-value was 1/301.

Effect sizes included the observed-minus-null differences in the number and fraction of significant cells, as well as the fold change in significant-cell fraction. A pathway was classified as non-randomly prominent only if it met all three prespecified criteria: a within-patient empirical FDR of ≤0.05, an observed-minus-null difference in significant-cell fraction of ≥0.02, and a Z score of ≥2.0.

## Data Availability

The single-cell RNA-seq data, gene annotations, cell-type labels, and sample metadata from the Wu et al. breast cancer cohort were accessed through 3CA^1^. The HIPPIE^42^ interactome and KEGG_2021_Human gene sets from Enrichr (https://maayanlab.cloud/Enrichr/) were used for network and pathway analyses.

## Code Availability

The code for the analyses can be accessed from the following repository: https://github.com/bengiruken

## Acknowledgements

This Research was supported by the Cancer Innovation Laboratory, Center for Cancer Research, National Cancer Institute, National Institutes of Health Intramural Research Program project numbers, ZIA BC 010441 and ZIA BC 010442, and federal funds from the National Cancer Institute, National Institutes of Health, under contract HHSN261201500003I. The contributions of the NIH authors were made as part of their official duties as NIH federal employees, are in compliance with agency policy requirements, and are considered Works of the United States Government. However, the findings and conclusions presented in this paper are those of the authors and do not necessarily reflect the views of the NIH or the U.S. Department of Health and Human Services.

## Author contributions

B.R.Y., H.J., and R.N. conceived and designed the study. B.R.Y. did the data curation and visualization, analyzed the results and drafted the manuscript. B.R.Y., H.J., and R.N. validated, reviewed, and edited the manuscript. R.N. supervised the project.

## References

1 Wu, S. Z. et al. A single-cell and spatially resolved atlas of human breast cancers. Nat Genet 53, 1334–1347 (2021). 10.1038/s41588-021-00911-1

2 Liu, S. Q. et al. Single-cell and spatially resolved analysis uncovers cell heterogeneity of breast cancer. J Hematol Oncol 15, 19 (2022). 10.1186/s13045-022-01236-0

3 Ozmen, F. et al. Single-cell RNA sequencing reveals different cellular states in malignant cells and the tumor microenvironment in primary and metastatic ER-positive breast cancer. NPJ Breast Cancer 11, 95 (2025). 10.1038/s41523-025-00808-w

4 Azizi, E. et al. Single-Cell Map of Diverse Immune Phenotypes in the Breast Tumor Microenvironment. Cell 174, 1293–1308 e1236 (2018). 10.1016/j.cell.2018.05.060

5 Bassez, A. et al. A single-cell map of intratumoral changes during anti-PD1 treatment of patients with breast cancer. Nat Med 27, 820–832 (2021). 10.1038/s41591-021-01323-8

6 Fan, J., Slowikowski, K. & Zhang, F. Single-cell transcriptomics in cancer: computational challenges and opportunities. Exp Mol Med 52, 1452–1465 (2020). 10.1038/s12276-020-0422-0

7 Regner, M. J. et al. Defining the regulatory logic of breast cancer using single-cell epigenetic and transcriptome profiling. Cell Genom 5, 100765 (2025). 10.1016/j.xgen.2025.100765

8 Aibar, S. et al. SCENIC: single-cell regulatory network inference and clustering. Nat Methods 14, 1083–1086 (2017). 10.1038/nmeth.4463

9 Xu, L. et al. A comprehensive single-cell breast tumor atlas defines epithelial and immune heterogeneity and interactions predicting anti-PD-1 therapy response. Cell Rep Med 5, 101511 (2024). 10.1016/j.xcrm.2024.101511

10 Ors, A. et al. Estrogen regulates divergent transcriptional and epigenetic cell states in breast cancer. Nucleic Acids Res 50, 11492–11508 (2022). 10.1093/nar/gkac908

11 Roda, N. et al. A Rare Subset of Primary Tumor Cells with Concomitant Hyperactivation of Extracellular Matrix Remodeling and dsRNA-IFN1 Signaling Metastasizes in Breast Cancer. Cancer Res 83, 2155–2170 (2023). 10.1158/0008-5472.CAN-22-2717

12 Kabeer, F. et al. Single-cell decoding of drug induced transcriptomic reprogramming in triple negative breast cancers. Genome Biol 25, 191 (2024). 10.1186/s13059-024-03318-3

13 Menyailo, M. E. et al. Heterogeneity of Circulating Epithelial Cells in Breast Cancer at Single-Cell Resolution: Identifying Tumor and Hybrid Cells. Adv Biol (Weinh) 7, e2200206 (2023). 10.1002/adbi.202200206

14 Orbach, S. M. et al. Single-cell RNA-sequencing identifies anti-cancer immune phenotypes in the early lung metastatic niche during breast cancer. Clin Exp Metastasis 39, 865–881 (2022). 10.1007/s10585-022-10185-4

15 Nussinov, R., Tsai, C. J. & Jang, H. Signaling in the crowded cell. Curr Opin Struct Biol 71, 43–50 (2021). 10.1016/j.sbi.2021.05.009

16 Yavuz, B. R., Tsai, C. J., Nussinov, R. & Tuncbag, N. Pan-cancer clinical impact of latent drivers from double mutations. Commun Biol 6, 202 (2023). 10.1038/s42003-023-04519-5

17 Nussinov, R., Yavuz, B. R. & Jang, H. Molecular principles underlying aggressive cancers. Signal Transduct Target Ther 10, 42 (2025). 10.1038/s41392-025-02129-7

18 Denisenko, T. V. et al. Signalomics for molecular tumor boards and precision oncology of breast and gynecological cancers. Mol Syst Biol 21, 952–959 (2025). 10.1038/s44320-025-00125-1

19 Alam, J., Huda, M. N., Tackett, A. J. & Miah, S. Oncogenic signaling-mediated regulation of chromatin during tumorigenesis. Cancer Metastasis Rev 42, 409–425 (2023). 10.1007/s10555-023-10104-3

20 Goenka, A. et al. Tumor microenvironment signaling and therapeutics in cancer progression. Cancer Commun (Lond) 43, 525–561 (2023). 10.1002/cac2.12416

21 Li, Y. J., Zhang, C., Martincuks, A., Herrmann, A. & Yu, H. STAT proteins in cancer: orchestration of metabolism. Nat Rev Cancer 23, 115–134 (2023). 10.1038/s41568-022-00537-3

22 Dakal, T. C. et al. Intricate relationship between cancer stemness, metastasis, and drug resistance. MedComm (2020) 5, e710 (2024). 10.1002/mco2.710

23 Ryspayeva, D. et al. Signaling pathway dysregulation in breast cancer. Oncotarget 16, 168–201 (2025). 10.18632/oncotarget.28701

24 Long, L., Fei, X., Chen, L., Yao, L. & Lei, X. Potential therapeutic targets of the JAK2/STAT3 signaling pathway in triple-negative breast cancer. Front Oncol 14, 1381251 (2024). 10.3389/fonc.2024.1381251

25 Han, M. et al. MulNet: a scalable framework for reconstructing intra- and intercellular signaling networks from bulk and single-cell RNA-seq data. Brief Bioinform 26 (2025). 10.1093/bib/bbaf081

26 Cha, J., Lavi, M., Kim, J., Shomron, N. & Lee, I. Imputation of single-cell transcriptome data enables the reconstruction of networks predictive of breast cancer metastasis. Comput Struct Biotechnol J 21, 2296–2304 (2023). 10.1016/j.csbj.2023.03.036

27 Feng, K. et al. Integrative analysis of bulk and single-cell transcriptomics identifies factors related to immunosuppressive microenvironment to predict unfavorable prognosis in inflammatory breast cancer. Front Immunol 16, 1727590 (2025). 10.3389/fimmu.2025.1727590

28 Gao, X. et al. The integrated single-cell analysis interpret the lactate metabolism-driven immune suppression in triple-negative breast cancer. Discov Oncol 16, 784 (2025). 10.1007/s12672-025-02605-0

29 Liu, F., Zhang, J., Gu, X., Guo, Q. & Guo, W. Single-cell transcriptome sequencing reveals SPP1-CD44-mediated macrophage-tumor cell interactions drive chemoresistance in TNBC. J Cell Mol Med 28, e18525 (2024). 10.1111/jcmm.18525

30 Zhang, Y. et al. Distinct cellular mechanisms underlie chemotherapies and PD-L1 blockade combinations in triple-negative breast cancer. Cancer Cell 43, 446–463 e447 (2025). 10.1016/j.ccell.2025.01.007

31 Feng, J. et al. sc2MeNetDrug: A computational tool to uncover inter-cell signaling targets and identify relevant drugs based on single cell RNA-seq data. PLoS Comput Biol 20, e1011785 (2024). 10.1371/journal.pcbi.1011785

32 Ianevski, A. et al. Single-cell transcriptomes identify patient-tailored therapies for selective co-inhibition of cancer clones. Nat Commun 15, 8579 (2024). 10.1038/s41467-024-52980-5

33. Osorio, D., Shahrouzi, P., Tekpli, X., Kristensen, V. N. & Kuijjer, M. L. (eLife Sciences Publications, Ltd, 2025).

34 Tang, C. et al. Personalized tumor combination therapy optimization using the single-cell transcriptome. Genome Med 15, 105 (2023). 10.1186/s13073-023-01256-6

35 Wang, X. et al. An integrated computational strategy to predict personalized cancer drug combinations by reversing drug resistance signatures. Comput Biol Med 163, 107230 (2023). 10.1016/j.compbiomed.2023.107230

36 You, T. et al. A highly annotated drug combination resource for catalyzing precision combinatorial therapy. Sci Data 12, 1284 (2025). 10.1038/s41597-025-05630-4

37 Vis, D. J. et al. A pan-cancer screen identifies drug combination benefit in cancer cell lines at the individual and population level. Cell Rep Med 5, 101687 (2024). 10.1016/j.xcrm.2024.101687

38 Tosh, C. et al. A Bayesian active learning platform for scalable combination drug screens. Nat Commun 16, 156 (2025). 10.1038/s41467-024-55287-7

39 Yavuz, B. R., Jang, H. & Nussinov, R. Discovering anticancer drug target combinations via network-informed signaling-based approach. Commun Med (Lond) 5, 428 (2025). 10.1038/s43856-025-01150-9

40 Nussinov, R., Yavuz, B. R. & Jang, H. Anticancer drugs: How to select small molecule combinations? Trends Pharmacol Sci 45, 503–519 (2024). 10.1016/j.tips.2024.04.012

41 Tyler, M. et al. The Curated Cancer Cell Atlas provides a comprehensive characterization of tumors at single-cell resolution. Nat Cancer 6, 1088–1101 (2025). 10.1038/s43018-025-00957-8

42 Alanis-Lobato, G., Andrade-Navarro, M. A. & Schaefer, M. H. HIPPIE v2.0: enhancing meaningfulness and reliability of protein-protein interaction networks. Nucleic Acids Res 45, D408–D414 (2017). 10.1093/nar/gkw985

43 Yavuz, B. R., Sahin, U., Jang, H., Nussinov, R. & Tuncbag, N. Mutations in tumor signaling, metastases, and synthetic lethality establish distinct patterns. PLoS Comput Biol 21, e1013351 (2025). 10.1371/journal.pcbi.1013351

44 Kanehisa, M. & Goto, S. KEGG: kyoto encyclopedia of genes and genomes. Nucleic Acids Res 28, 27–30 (2000). 10.1093/nar/28.1.27

45 Maxwell, P. H. et al. The tumour suppressor protein VHL targets hypoxia-inducible factors for oxygen-dependent proteolysis. Nature 399, 271–275 (1999). 10.1038/20459

46 Baek, J. H. et al. OS-9 interacts with hypoxia-inducible factor 1alpha and prolyl hydroxylases to promote oxygen-dependent degradation of HIF-1alpha. Mol Cell 17, 503–512 (2005). 10.1016/j.molcel.2005.01.011

47 Semenza, G. L. Defining the role of hypoxia-inducible factor 1 in cancer biology and therapeutics. Oncogene 29, 625–634 (2010). 10.1038/onc.2009.441

48 Singh, D. et al. Overexpression of hypoxia-inducible factor and metabolic pathways: possible targets of cancer. Cell Biosci 7, 62 (2017). 10.1186/s13578-017-0190-2

49 Gilkes, D. M. & Semenza, G. L. Role of hypoxia-inducible factors in breast cancer metastasis. Future Oncol 9, 1623–1636 (2013). 10.2217/fon.13.92

50 Zheng, X. D., Li, H. Y., Gao, S. Y., Wang, Q. & Liu, J. B. High hypoxia inducible factor-1alpha expression is associated with reduced survival in patients with breast cancer: A meta-analysis. World J Clin Oncol 16, 105691 (2025). 10.5306/wjco.v16.i6.105691

51 Zhi, S. et al. Hypoxia-inducible factor in breast cancer: role and target for breast cancer treatment. Front Immunol 15, 1370800 (2024). 10.3389/fimmu.2024.1370800

52 Shi, Y. & Gilkes, D. M. HIF-1 and HIF-2 in cancer: structure, regulation, and therapeutic prospects. Cell Mol Life Sci 82, 44 (2025). 10.1007/s00018-024-05537-0

53 He, W., Batty-Stuart, S., Lee, J. E. & Ohh, M. HIF-1alpha Hydroxyprolines Modulate Oxygen-Dependent Protein Stability Via Single VHL Interface With Comparable Effect on Ubiquitination Rate. J Mol Biol 433, 167244 (2021). 10.1016/j.jmb.2021.167244

54 Zhang, J., Yao, M., Xia, S., Zeng, F. & Liu, Q. Systematic and comprehensive insights into HIF-1 stabilization under normoxic conditions: implications for cellular adaptation and therapeutic strategies in cancer. Cell Mol Biol Lett 30, 2 (2025). 10.1186/s11658-024-00682-7

55 Laughner, E., Taghavi, P., Chiles, K., Mahon, P. C. & Semenza, G. L. HER2 (neu) signaling increases the rate of hypoxia-inducible factor 1alpha (HIF-1alpha) synthesis: novel mechanism for HIF-1-mediated vascular endothelial growth factor expression. Mol Cell Biol 21, 3995–4004 (2001). 10.1128/MCB.21.12.3995-4004.2001

56 Dodd, K. M., Yang, J., Shen, M. H., Sampson, J. R. & Tee, A. R. mTORC1 drives HIF-1alpha and VEGF-A signalling via multiple mechanisms involving 4E-BP1, S6K1 and STAT3. Oncogene 34, 2239–2250 (2015). 10.1038/onc.2014.164

57 Pore, N. et al. Akt1 activation can augment hypoxia-inducible factor-1alpha expression by increasing protein translation through a mammalian target of rapamycin-independent pathway. Mol Cancer Res 4, 471–479 (2006). 10.1158/1541-7786.MCR-05-0234

58 Gort, E. H. et al. Hypoxia-inducible factor-1alpha expression requires PI 3-kinase activity and correlates with Akt1 phosphorylation in invasive breast carcinomas. Oncogene 25, 6123–6127 (2006). 10.1038/sj.onc.1209643

59 Nussinov, R., Tsai, C. J. & Jang, H. Does Ras Activate Raf and PI3K Allosterically? Front Oncol 9, 1231 (2019). 10.3389/fonc.2019.01231

60 Sang, N. et al. MAPK signaling up-regulates the activity of hypoxia-inducible factors by its effects on p300. J Biol Chem 278, 14013–14019 (2003). 10.1074/jbc.M209702200

61 Mylonis, I. et al. Identification of MAPK phosphorylation sites and their role in the localization and activity of hypoxia-inducible factor-1alpha. J Biol Chem 281, 33095– 33106 (2006). 10.1074/jbc.M605058200

62 Sodhi, A., Montaner, S., Miyazaki, H. & Gutkind, J. S. MAPK and Akt act cooperatively but independently on hypoxia inducible factor-1alpha in rasV12 upregulation of VEGF. Biochem Biophys Res Commun 287, 292–300 (2001). 10.1006/bbrc.2001.5532

63 Xu, Q. et al. Targeting Stat3 blocks both HIF-1 and VEGF expression induced by multiple oncogenic growth signaling pathways. Oncogene 24, 5552–5560 (2005). 10.1038/sj.onc.1208719

64 Niu, G. et al. Signal transducer and activator of transcription 3 is required for hypoxia-inducible factor-1alpha RNA expression in both tumor cells and tumor-associated myeloid cells. Mol Cancer Res 6, 1099–1105 (2008). 10.1158/1541-7786.MCR-07-2177

65 Jung, J. E. et al. STAT3 inhibits the degradation of HIF-1alpha by pVHL-mediated ubiquitination. Exp Mol Med 40, 479–485 (2008). 10.3858/emm.2008.40.5.479

66 Pawlus, M. R., Wang, L. & Hu, C. J. STAT3 and HIF1alpha cooperatively activate HIF1 target genes in MDA-MB-231 and RCC4 cells. Oncogene 33, 1670–1679 (2014). 10.1038/onc.2013.115

67 Dinarello, A. et al. STAT3 and HIF1alpha cooperatively mediate the transcriptional and physiological responses to hypoxia. Cell Death Discov 9, 226 (2023). 10.1038/s41420-023-01507-w

68 Zhao, F. L. & Qin, C. F. EGF promotes HIF-1alpha expression in colorectal cancer cells and tumor metastasis by regulating phosphorylation of STAT3. Eur Rev Med Pharmacol Sci 23, 1055–1062 (2019). 10.26355/eurrev_201902_16993

69 Shaikat, A. H. et al. Investigating hypoxia-inducible factor signaling in cancer: Mechanisms, clinical implications, targeted therapeutic strategies, and resistance. Cancer Pathog Ther 4, 174–191 (2026). 10.1016/j.cpt.2025.07.003

70 Zhang, Z., Yao, L., Yang, J., Wang, Z. & Du, G. PI3K/Akt and HIF-1 signaling pathway in hypoxia-ischemia (Review). Mol Med Rep 18, 3547–3554 (2018). 10.3892/mmr.2018.9375

71 Tian, Y. et al. PI3K/AKT signaling activates HIF1alpha to modulate the biological effects of invasive breast cancer with microcalcification. NPJ Breast Cancer 9, 93 (2023). 10.1038/s41523-023-00598-z

72 Courtnay, R. et al. Cancer metabolism and the Warburg effect: the role of HIF-1 and PI3K. Mol Biol Rep 42, 841–851 (2015). 10.1007/s11033-015-3858-x

73 Balamurugan, K. et al. C/EBPdelta links IL-6 and HIF-1 signaling to promote breast cancer stem cell-associated phenotypes. Oncogene 38, 3765–3780 (2019). 10.1038/s41388-018-0516-5

74 Su, Q. et al. Sanguinarine disrupts the colocalization and interaction of HIF-1alpha with tyrosine and serine phosphorylated-STAT3 in breast cancer. J Cell Mol Med 24, 3756–3761 (2020). 10.1111/jcmm.15056

75 Smith, A. E. et al. Tipifarnib Potentiates the Antitumor Effects of PI3Kalpha Inhibition in PIK3CA- and HRAS-Dysregulated HNSCC via Convergent Inhibition of mTOR Activity. Cancer Res 83, 3252–3263 (2023). 10.1158/0008-5472.CAN-23-0282

76 Algazi, A. P. et al. A phase 1 study of triple-targeted therapy with BRAF, MEK, and AKT inhibitors for patients with BRAF-mutated cancers. Cancer 130, 1784–1796 (2024). 10.1002/cncr.35200

77 Turner, N. C. et al. Inavolisib-Based Therapy in PIK3CA-Mutated Advanced Breast Cancer. N Engl J Med 391, 1584–1596 (2024). 10.1056/NEJMoa2404625

78 Andre, F. et al. Alpelisib for PIK3CA-Mutated, Hormone Receptor-Positive Advanced Breast Cancer. N Engl J Med 380, 1929–1940 (2019). 10.1056/NEJMoa1813904

79 Jhaveri, K. L. et al. Overall Survival with Inavolisib in PIK3CA-Mutated Advanced Breast Cancer. N Engl J Med 393, 151–161 (2025). 10.1056/NEJMoa2501796

80 Sacher, A. et al. Single-Agent Divarasib (GDC-6036) in Solid Tumors with a KRAS G12C Mutation. N Engl J Med 389, 710–721 (2023). 10.1056/NEJMoa2303810

81 Murciano-Goroff, Y. R. et al. Pan-tumor activity of olomorasib, a next-generation KRAS G12C inhibitor in KRAS G12C-mutant advanced solid tumors: a first-in-human study. Nat Commun 17 (2026). 10.1038/s41467-026-69943-7

82 Wolpin, B. M. et al. Daraxonrasib in Previously Treated Advanced RAS-Mutated Pancreatic Cancer. N Engl J Med 394, 1790–1802 (2026). 10.1056/NEJMoa2505783

83 Wolf, F. A., Angerer, P. & Theis, F. J. SCANPY: large-scale single-cell gene expression data analysis. Genome Biol 19, 15 (2018). 10.1186/s13059-017-1382-0

84 Langville, A. N. & Meyer, C. D. A Survey of Eigenvector Methods for Web Information Retrieval. SIAM Review 47, 135–161 (2005). 10.1137/s0036144503424786

85 Brin, S. & Page, L. The anatomy of a large-scale hypertextual web search engine (Reprint from COMPUTER NETWORKS AND ISDN SYSTEMS, vol 30, pg 107–117, 1998). Computer networks 56, 3825–3833 (2012).

86 Kuleshov, M. V. et al. Enrichr: a comprehensive gene set enrichment analysis web server 2016 update. Nucleic Acids Res 44, W90–97 (2016). 10.1093/nar/gkw377

87 North, B. V., Curtis, D. & Sham, P. C. A note on the calculation of empirical P values from Monte Carlo procedures. Am J Hum Genet 71, 439–441 (2002). 10.1086/341527

88 Phipson, B. & Smyth, G. K. Permutation P-values should never be zero: calculating exact P-values when permutations are randomly drawn. Stat Appl Genet Mol Biol 9, Article39 (2010). 10.2202/1544-6115.1585

