## Supplementary Figures for "Single-Cell Mapping of Malignant Signaling Networks Guides Drug Combinations"

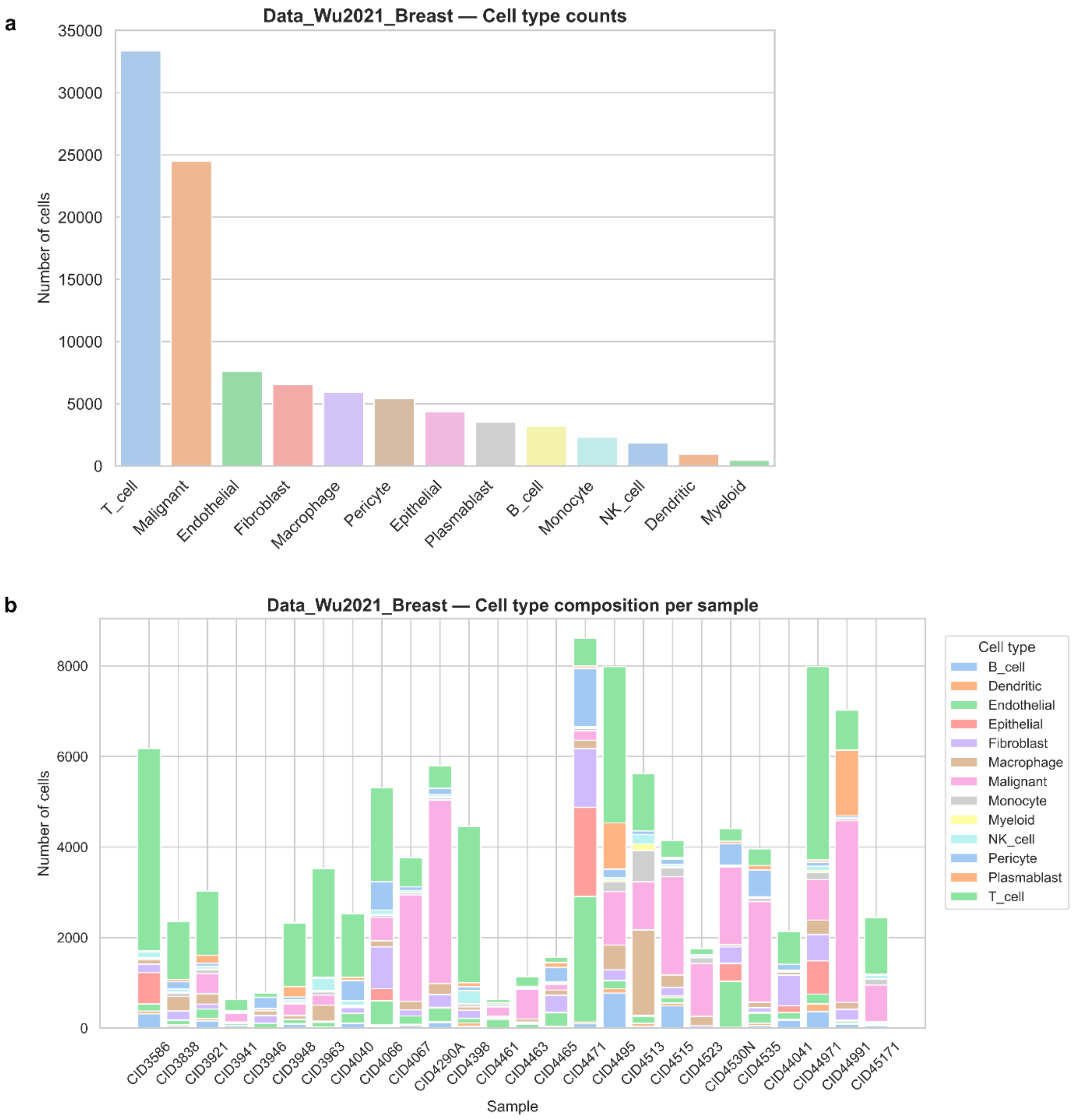


**Figure S1. (a)** The breast cancer dataset from Wu et al., contains 100,064 annotated cells across 13 cell types. T cells are the most abundant population, with 33,368 cells, followed by malignant cells with 24,489 cells. Other represented cell types included endothelial cells, fibroblasts, macrophages, pericytes, epithelial cells, plasmablasts, B cells, monocytes, NK cells, dendritic cells, and myeloid cells, with counts ranging from 7,605 endothelial cells to 463 myeloid cells. **(b)** The per-sample stacked bar plot shows that both total cell number and cell-type composition varied across samples.

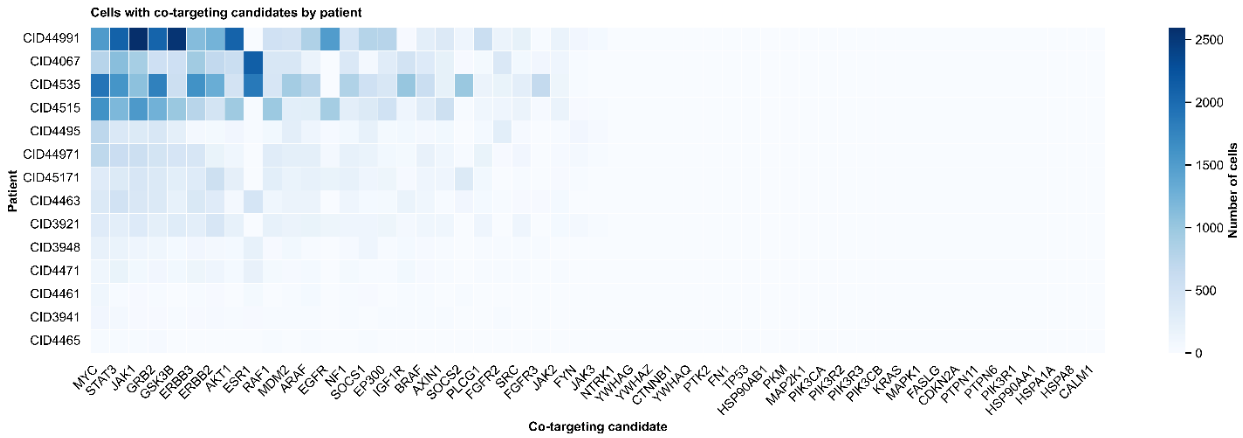


**Figure S2. Patient-wise distribution of co-targeting candidate cell counts.** Heatmap showing the number of malignant treatment-naïve cells associated with each co-targeting candidate gene across patients. Rows represent individual patients, and columns represent co-targeting candidate genes. For each patient–gene pair, the color intensity indicates the number of cells in which that candidate was identified, with darker blue corresponding to higher cell counts and near-white corresponding to low or zero counts. The color bar indicates the absolute number of cells. Patient-level differences should be interpreted together with the total malignant-cell representation per patient, because patients with more retained malignant cells may contribute higher absolute candidate counts.

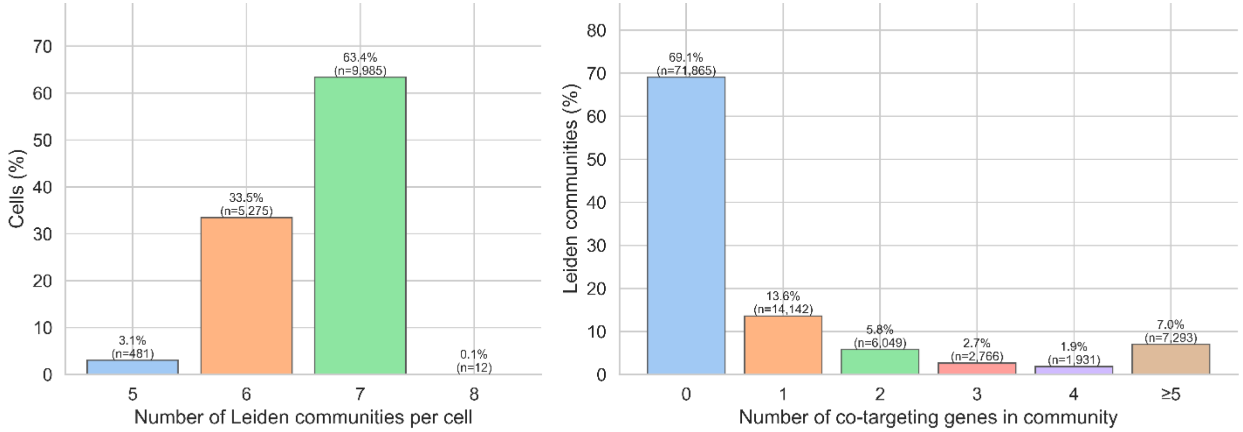

**Figure S3. Distribution of Leiden community number per cell and co-targeting gene content per community. (left)** Percentage of malignant treatment-naïve cells containing each number of Leiden communities. Bars show the proportion of cells with 5, 6, 7, or 8 Leiden communities, with labels indicating both percentage and cell count. Most cells contained 7 Leiden communities, followed by 6 communities. **(right)** Percentage of Leiden communities containing 0, 1, 2, 3, 4, or ≥5 co-targeting candidate genes. Bars show the proportion of Leiden communities in each co-targeting gene-count category, with labels indicating both percentage and number of communities. Co-targeting gene counts were collapsed at ≥5 to summarize the long-tailed distribution. Most Leiden communities contained no co-targeting genes, while a smaller subset contained one or more co-targeting candidates.

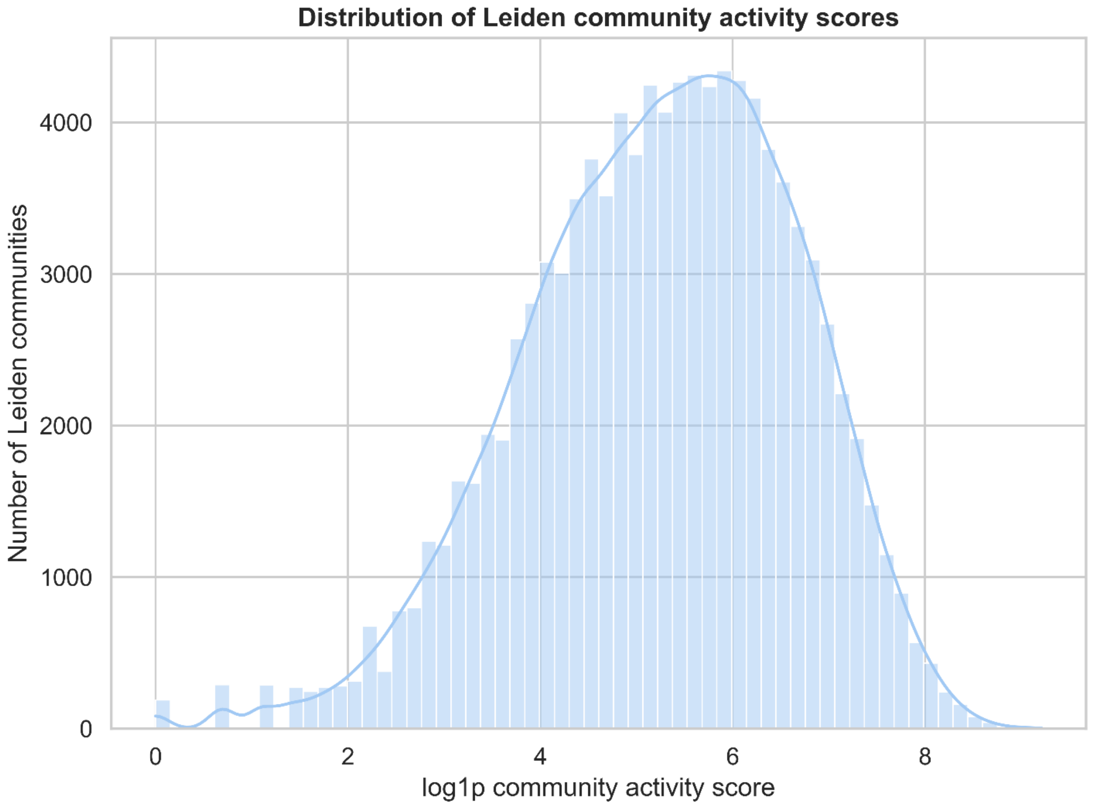


**Figure S4. Distribution of Leiden community activity scores and selected high-scoring communities across patients.** Histogram showing the overall distribution of log1p-transformed values across cell-specific Leiden communities. The x-axis shows the log1p-transformed community activity score, and the y-axis shows the number of Leiden communities. This panel summarizes the range of aggregate expression-based activity scores observed across Leiden communities before selecting the highest-scoring community within each cell.

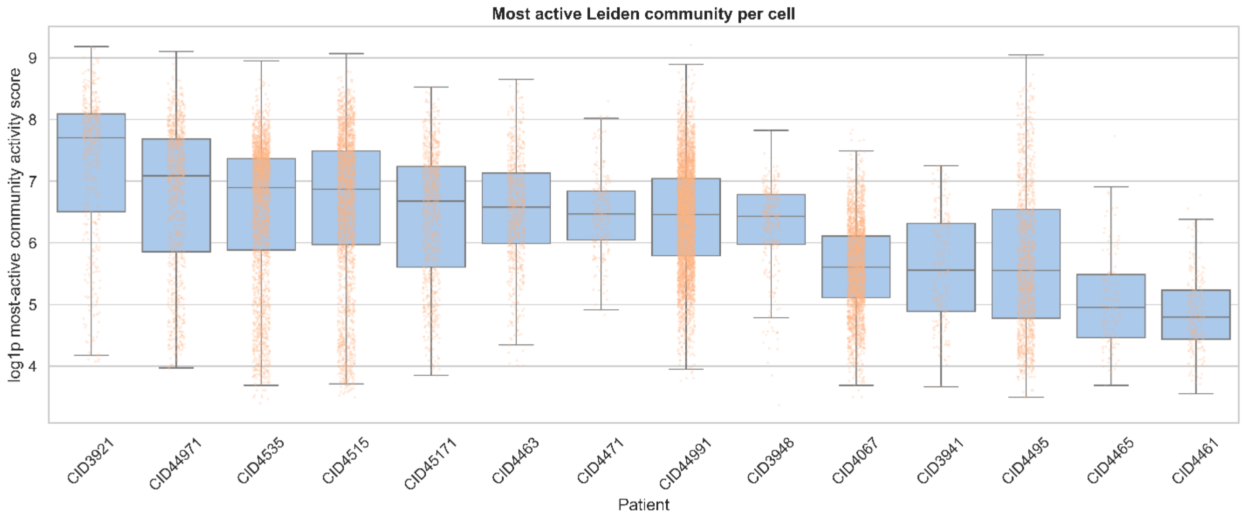


**Figure S5. Patient-level distribution of selected highest-scoring Leiden community activity scores.** For each malignant untreated cell, the Leiden community with the highest precomputed CommunityActivityScore was selected as the top-ranked within-cell module, with the number of genes in the community used as an additional ranking criterion when needed. Each point represents one cell, and boxplots summarize patient-level distributions of log1p-transformed selected community activity scores. The plot summarizes both patient-level differences and cell-to-cell variability in aggregate expression-based scores of the selected Leiden communities.
